# The molecular mass and isoelectric point of plant proteomes

**DOI:** 10.1101/546077

**Authors:** Tapan Kumar Mohanta, Abdullatif Khan, Abeer Hashem, Elsayed Fathi Abd_Allah, Ahmed Al-Harrasi

**Affiliations:** Natural and Medical Science Research Centre, University of Nizwa, 616 Nizwa, Oman; Botany and Microbiology Department, King Saud University, Riyadh 11451, Saudi Arabia; Plant Production Department, King Saud University, Riyadh 11451, Saudi Arabia

**Keywords:** Proteome, Amino acids, Isoelectric point, Molecular weight, Selenocysteine, Pyrrolysine

## Abstract

**Background:** Cell contain diverse array of proteins with different molecular weight and isoelectric point (pI). The molecular weight and pI of protein play important role in determining the molecular biochemical function. Therefore, it was important to understand the detail regarding the molecular weight and pI of the plant proteins.

**Results:** A proteome-wide analysis of plant proteomes from 145 species revealed a *pI* range of 1.99 (epsin) to 13.96 (hypothetical protein). The spectrum of molecular mass of the plant proteins varied from 0.54 to 2236.8 kDa. A putative Type-I polyketide synthase (22244 amino acids) in *Volvox carteri* was found to be the largest protein in the plant kingdom. However, Type-I polyketide synthase was not found in higher plant species. Titin (806.46 kDa) and misin/midasin (730.02 kDa) were the largest proteins identified in higher plant species. The *pI* and molecular weight of the plant proteins showed a trimodal distribution. An acidic *pI* (56.44% of proteins) was found to be predominant over a basic *pI* (43.34% of proteins) and the abundance of acidic *pI* proteins was higher in unicellular algae species relative to multicellular higher plants. In contrast, the seaweed, *Porphyra umbilicalis*, possesses a higher proportion of basic *pI* proteins (70.09%). Plant proteomes were also found to contain selenocysteine (Sec), amino acid that was found only in lower eukaryotic aquatic plant lineage. Amino acid composition analysis showed Leu was high and Trp was low abundant amino acids in the plant proteome. Additionally, the plant proteomes also possess ambiguous amino acids Xaa (unknown), Asx (asparagine or aspartic acid), Glx (glutamine or glutamic acid), and Xle (leucine or isoleucine) as well.

**Conclusion:** The diverse molecular weight and isoelectric point range of plant proteome will be helpful to understand their biochemical and functional aspects. The presence of selenocysteine proteins in lower eukaryotic organism is of interest and their expression in higher plant system can help us to understand their functional role.

## Background

The isoelectric or isoionic point of a protein is the pH at which a protein carries no net electrical charge and hence is considered neutral [1–4]. The zwitterion form of a protein becomes dominant at neutral pH. The *pI* of polypeptides is largely dependent on the dissociation constant of the ionisable groups [5]. The major ionisable groups present in the amino acids are arginine, aspartate, cysteine, histidine, glutamate, lysine, and glutamate, where they play a major role in determining the *pI* of a protein [6–8]. Co-translational and post-translational modifications of a protein, however, can also play a significant role in determining the *pI* of a protein [9, 10]. The exposure of charged residues to the solvents, hydrogen bonds (diploe interactions) and dehydration also impact the *pI* of a protein [11, 12]. The inherent *pI* of protein, however, is primarily based on its native protein sequence. The *pI* of a protein is crucial to understanding its biochemical function and thus determining *pI* is an essential aspect of proteomic studies. During electrophoresis, the direction of movement of a protein in a gel or other matrix depends its’*pI*, hence numerous proteins can be separated based on their *pI* [13–16]. Given the impact of post-translational modifications and other biochemical alterations (phosphorylation, methylation, alkylation), however, the predicted *pI* of a protein will certainly be different than the predicted *pI*; the latter of which is based on the composition of amino acids in a protein [9, 17, 18]. Nonetheless, an estimated isoelectric point is highly important and a commonly identified parameter.

Several studies have been conducted to understand the *pI* of proteins/polypeptides [3, 19–21]. These studies have been mainly based on animal, bacteria, and virus models and databases containing the *pI* of experimentally verified proteins. None of these databases, however, contain more than ten thousand proteins sequences which is very few relative to the availability of proteomic data. Therefore, an analysis was conducted of the *pI* and molecular weight of proteins from 144 plant species which included 5.87 million protein sequences. This analysis provides an in-depth analysis of the *pI* and molecular mass of the proteins in the plant kingdom.

## Results and discussion

### Plant proteins range from 0.54 kDa to 2236.8 kDa

A proteome-based analysis of plant proteins of 144 plant species that included more than 5.86 million protein sequences was considered to study the molecular mass, *pI*, and amino acid composition of proteins that exist in plant proteomes (Additional file 2: Table S1). The analysis indicated that *Hordeum vulgare* possessed the highest number (248180) of protein sequences, while *Helicosporidium* sp. had the lowest number (6033). On average, plant proteomes possess 40988.66 protein sequences per species (Fig. 1). The analysis also revealed that the molecular mass of plant proteomes ranged from 0.54 kDa to 2236.8 kDa. The gross molecular weight of plant proteome was ranged from 9857470.16 kDa (*Hordeum vulgare*) to 178551.42 kDa (*Helicosporidium sp.*) (Additional file Table S1). *Volvox carteri* was found to possess the largest plant protein (XP_002951836.1) of 2236.8 kDa, containing 22244 amino acids (*pI* 5.94), while *Citrus unshiu* possessed the smallest protein of 0.54 kDa, containing only four amino acids (*pI* 5.98) (id: GAY42954.1). This is the first analysis to document the largest (2236.8 kDa) and smallest (0.54 kDa) protein in the plant kingdom. These smallest and largest proteins yet to be functionally annotated. The BLASTP analysis in the NCBI database did not identify suitable similarity with any other proteins. Some of the domains identified in the largest protein were matched with protein domains of Type-I polyketide synthase. The molecular mass of some other high molecular mass proteins were: 2056.44 kDa (id: XP_001698501.1, type-1 polyketide synthase, *pI*: 6.00, aa: 21004, *Chlamydomonas reinhardtii*); 1994.71 kDa (id: XP_ 001416378.1, polyketide synthase, *pI*: 7.38, aa: 18193, *Ostreococcus lucimarinus*); 1932.21 kDa (id: Cz02g22160.t1, unknown protein, *pI*: 5.7, aa: 18533, *Chromochloris zofingiensis*); 1814.1 kDa (id: XP_007509537.1, unknown protein, *pI*: 4.46, aa: 16310, *Bathycoccus prasinos*); 1649.26 kDa (id: XP_011401890.1, polyketide synthase, *pI*: 5.53, aa: 16440, *Auxenochlorella protothecoides*); 1632.35 kDa (id: XP_005650993.1, ketoacyl-synt-domain-containing protein, *pI*: 5.86, aa: 15797,*Coccomyxa subellipsoidea*); 1532.91 kDa (id: XP_002507643.1, polyketide synthase, *pI*: 7.07, aa: 14149, *Micromonas commoda*); 1370.23 kDa (id: GAX78753.1, hypothetical protein CEUSTIGMA, *pI*: 5.97, aa: 13200, *Chlamydomonas eustigma*); 1300.83 kDa (id: XP_022026115.1, unknown protein/filaggrin-like, *pI*: 11.75, aa: 12581, *Helianthus annuus*); 1269.42 kDa (id: XP_009350379.1, unknown protein, *pI*: 5.37, aa: 11880, *Pyrus bretschneideri*); 1237.34 kDa (id: XP_ 022840687.1, polyketide synthase, *pI*: 7.30, aa: 11265, *Ostreococcus tauri*); 1159.35 kDa (id: XP_005847912.1, polyketide synthase, *pI*: 5.91, aa: 11464, *Chlorella variabilis*); 1150.02 kDa (id: PKI66547.1, unknown protein, *pI*: 3.87, aa: 11234, *Punica granatum*); 1027.64 kDa (id: Sphfalx0133s0012.1, unknown protein, *pI*: 4.05, aa: 9126, *Sphagnum fallax*); 909.93 kDa (id: XP_002985373.1, unknown/titin-like protein, *pI*: 4.02, aa: 8462, *Selaginella moellendorffii*); 881.59 kDa (id: KXZ46216.1, hypothetical protein, *pI*: 5.80, aa: 8881, *Gonium pectorale*); 848.29 kDa (id: XP_003056330.1, *pI*: 6.12, aa: 7926, *Micromonas pusilla*); 813.31 kDa (id: GAQ82263.1, unknown protein, *pI*: 4.60, aa: 7617, *Klebsormidium nitens*), 806.46 kDa (id: XP_017639830.1, titin-like, *pI*: 4.21, aa: 7209, *Gossypium arboreum*); 806.12 kDa (id: OAE35580.1, *pI*: 4.83, hypothetical protein, aa: 7651, *Marchantia polymorpha*); and 802.74 kDa, (id: XP_012444755.1, titin-like, *pI*: 4.19, aa: 7181, *Gossypium raimondii*) (Additional file 2: Table S1).

**Fig. 1.**
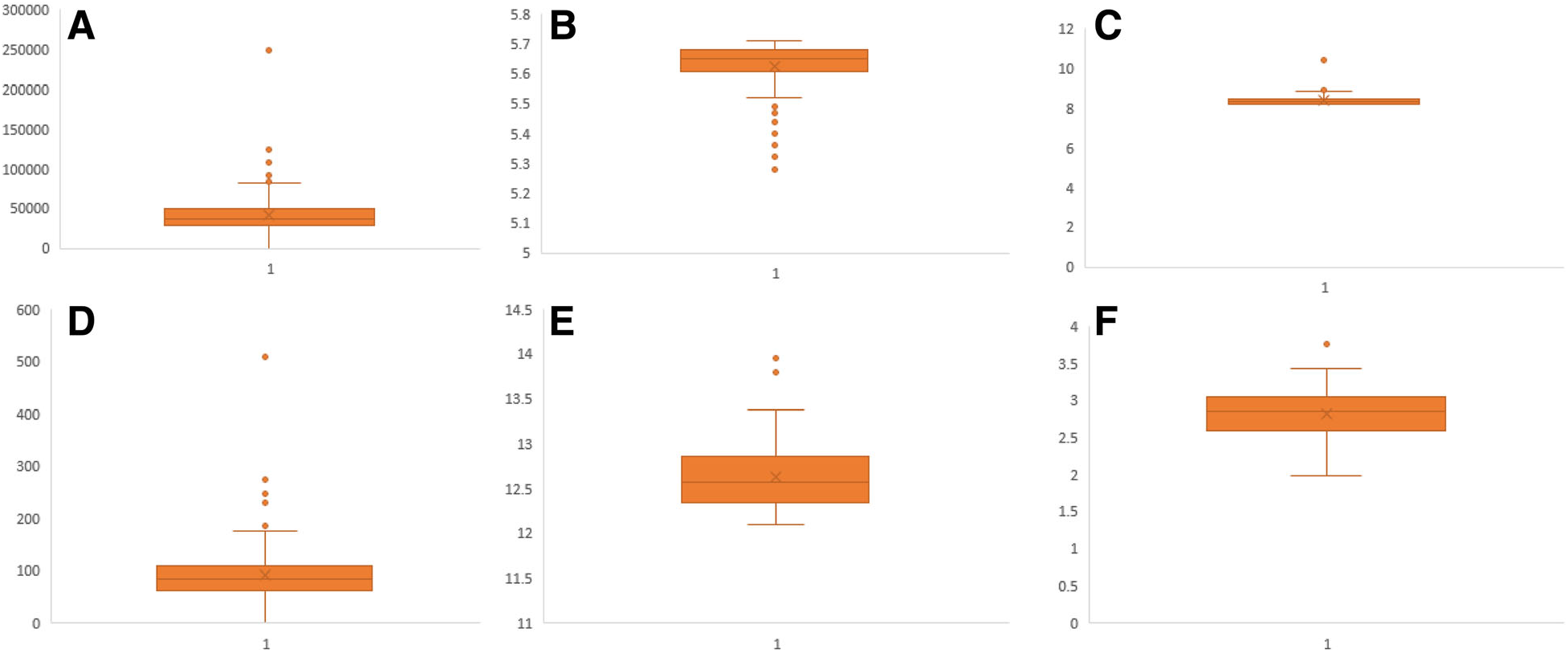
Box and Whisker plot of (**a**) average number of protein sequences encoded in plant proteome, **b** average acidic pI, **c** average basic pI of plant proteome, **d** average number of neutral pI proteins per plant proteome (**e**) average of highest pI protein, and (**f**) average of lowest pI proteins. Details can be found in Additional file 2: Table S1

On average, approximately 7.38% of the analysed proteins were found to contain ≥100 kDa proteins. Analysis of average molecular weight of proteins in a proteome revealed, the average molecular weight of individual protein of *Chlamydomonas reinhardtii* proteome was found to be 62.74 kDa, followed by 60.24 kDa (*Volvox carteri*), 58.64 kDa (*Klebsmordium nitens*), and 58.10 kDa (*Bathycoccus prasinos*). Similarly, *Trifolium pratense* (28.37 kDa) encode the lowest molecular weight protein in the plant kingdom. The algal species, *V. carteri,* was found to encode largest plant protein (putative polyketide synthase); while other unicellular algae, and multi-cellular lower eukaryotic plants, including bryophytes and pteridophytes, were also found to encode some of the larger proteins (e.g. ketoacyl synthase) in the plant kingdom. The higher eukaryotic plants, including gymnosperms and angiosperms, were not found to encode a high molecular mass polyketide synthase protein. Notwithstanding, they contain the high molecular mass proteins; titin (806.46 kDa), misin/midasin (730.02 kDa), futsch (622.14 kDa), filaggrin (644.4 kDa), auxin transport protein BIG (568.4 kDa), and von Willebrand factor (624.74 kDa) (Additional file 2: Table S1). Titin protein found in human striated muscle is a large protein that is higher than 1 μM in length [22, 23]. The largest titin protein found in plants, however, was only 806.46 kDa (*Gossypium arboreum*). The predicted formula of the 806.46 kDa titin protein was C_33863_H_54610_N_9232_O_13061_S_200_ and its estimated half-life was 10–30 h; whereas the predicted formula of the 2236.8 kDa protein of *V. carteri* was C_97783_H_157401_N_28489_O_30265_S_637_. Almost all of the higher eukaryotic plants were found to possess titin, misin/midasin, and auxin transport protein BIG proteins. Species of unicellular algae were not found to possess titin or misin/ midasin proteins. This suggests that titin and misin/midasin proteins originated and evolved in more complex, multicellular organisms rather than unicellular organisms. Thus, the evolution of titin, misin/midasin proteins may also be associated with the evolution of terrestrial plants from aquatic plants. The Midasin protein.

The presence of the low molecular weight protein, other than the tripeptide glutathione (Cys, Gly, and Glu), was also determined. A 0.54 kDa molecular mass protein, containing only four amino acids (MIMF) and starting with methionine and ending with phenylalanine, was identified in *Citrus unshiu* (id: GAY42954.1) (Additional file 2: Table S1). Other low molecular mass plant proteins were 0.57 kDa (NP_001336532.1/ AT5G23115, *Arabidopsis thaliana*) and 0.63 kDa (AH003201-RA, *Amaranthus hypochondriacus*). Small proteins found in *A. thaliana* was MNPKS and that found in *A. hypochondriacus* was MLPYN, contained only five amino acids. The above mentioned low molecular mass proteins were missing in all of the studied species and their cellular and molecular functions have not been reported yet. One of the universal small molecular weight plant proteins, however, was identified as cytochrome b6/f complex subunit VIII (chloroplast) (MDIVSLAWAALM VVFTFSLSLVVWGRSGL) that contains only 29 amino acids. Cytochrome b6/f play important role in electron transfer system of photosystem II and administer photosynthesis [24–28]. It is commonly known that glutathione is the smallest functional polypeptide and that it plays diverse roles in cell signaling [29–31]. The tetra and penta peptides identified in the present analysis, however, were quite different from glutathione and none of them contained Cys, Gly, or Glu amino acids, as found in glutathione. Polypeptides with less than 100 amino acids are considered small proteins and studies indicate that considerable number of the small proteins are involved in cell signaling, DNA damage response, and cell growth [32–35]. During genome annotation, small protein-coding genes are overlooked constantly and hence they get buried amongst an enormous number of open reading frames [36]. Therefore, it is strenuous to find more numbers of small proteins in plants. The tetra and pentapeptides are involved in diverse signaling process [37, 38]. A bioactive tetrapeptide GEKG reported to boost extracellular matrix formation [39]. In bacteria, small peptides are act as pheromones [40]. The pentapeptide ERGMT regulate genetic competence in *B. subtilis* whereas ARNQT involved in sporulation [40].

A previously conducted comparative study revealed that plant proteins are comparatively smaller than animal proteins, as the former are encoded by fewer exons [41]. Longer proteins harbour more conserved domains and hence display a greater number of biological functions than short proteins. The average protein length of the studied plant species was 424.34 amino acids. A previous study reported the average length of eukaryotic proteins to be 472 amino acids and that the average length of plant proteins is approximately 81 amino acids shorter than animal proteins [41]. Our analysis indicates, however, that plant proteins are approximately 47.66 amino acid shorter than animal proteins. In addition, studies have also indicated that eukaryotic proteins are longer than bacterial proteins and that eukaryote genomes contain approximately 7 fold more proteins (48% larger) than bacterial genomes [42]. Although the average size of plant proteins was found to be 424.34 amino acids, the average protein size of lower, eukaryotic unicellular aquatic plant species; including *Chlamydomonas eustigma*, *Volvox carteri*, *Klebsordium nitens*, *Bathycoccus prasinos*, and *Durio zibethinus,* was found to be 576.56, 568.22, 538.73, 521.05, and 504.36 amino acids, respectively. This indicates that protein size unicellular plant species is larger than terrestrial multicellular complex plant species, suggesting that the evolution of plant proteins involved a loss of protein size and hence gene size. The reason for the alteration in protein length in the phylogenetic lineage of eukaryotic plants has yet to be elucidated. A multitude of evolutionary factors, including deletion (loss of exons) or fusion of multiple domains of proteins, most possibly played important roles in configuring the size of higher plant proteins. The insertion of transposon and splitting of genes expand the number of proteins but scale down the average size of the proteins [43–46]. Higher plants contain a very large number of transposable elements and therefore these elements are the most responsible factor to expect to have played a major role in increasing protein numbers and reducing the protein size in higher plants. The percentage of transposable elements in a genome is directly proportional to the genome size of the organism and varies from approximately 3% in small genomes to approximately 85% in large genomes [45]. Kirag et al. (2007) reported a significant correlation between protein length and the *pI* of a protein [19]. In our analysis, however, no correlation was found between protein length and the *pI* of a protein. For example, titin and misin are two of the larger proteins in the plants and they fall in the acidic *pI* range, but not in the alkaline *pI* range. Lack of sufficient proteomic data might be a possible factor for such results by Kirag et al., (2007).

### Plant encode a higher number of proteins than animals and fungi

Our analysis identified an average of 40469.47 proteins per genome (Additional file 2: Table S1). Previously the number of proteins in plant species was reported as 36795 per genome [41]. On average, animals and fungi encode 25189 and 9113 proteins per genome, respectively [41]. An average of 40469.47 proteins per plant genome is 62.24% higher than in animals and 444.08% higher than in fungi. Although, plant species encode a higher number of proteins, their size is smaller than the average size of animal proteins. Notably, green algae contain a smaller number of proteins than higher plants but their average protein size is 1.27 times larger. The average protein size (low to high) in the species of green algae ranged from 273.08 (*Helicosporidium* sp.) to 576.56 (*Chlamydomonas eustigma*) amino acids, dicots ranged from 253.34 (*Trifolium pratense*) to 498.49 (*Vitis vinifera*), and monocots ranged from 111.54 (*Hordeum vulgare*) to 473.35 (*Brachypodium distachyon*) amino acids. The average protein size of monocot proteins (431.07 amino acids), however, is slightly larger than dicots (424.3 amino acids). In addition to transposons, previous studies have reported that endosymbiosis may have also played an important role in the reduction of protein size in plant genomes [41, 47, 48]. This would have been due to the post endosymbiosis acquisition of thousands of genes from the chloroplast, since cyanobacterial proteins are smaller than eukaryotic proteins and cyanobacteria are the ancestors of plastids [41, 49]. In this hypothesis, the intermediate size of plant proteins would be the result of the migration of proteins from cyanobacteria (chloroplast) to the plant nucleus, thereby reducing the overall average size of the protein by a dilution effect [50, 51].

### The pI of plant proteins ranges from 1.99 to 13.96

Results indicated that the *pI* of analysed plant proteins ranged from 1.99 (id: PHT45033.1, *Capsicum baccatum*) to 13.96 (id: PKI59361.1, *Punica granatum*). The protein with the lowest *pI* (1.99) was epsin and the protein with the highest *pI* (13.96) was a hypothetical protein. We are the first to report on the plant proteins with the lowest and highest *pI*. The *C. baccatum* protein with *pI* 1.99 contains 271 amino acids, whereas the *P. granatum* protein with *pI* 13.96 contains 986 amino acids. The epsin protein (*pI* 1.99) is composed of 16 amino acid repeats (GWIDGWIDGWIDGW), while the hypothetical protein (*pI* 13.06) is composed of 64 QKLKSGLT and 31 TRRGLTAV repeats. From among the 20 essential amino acids, the epsin protein only contained five amino acids, namely Asp (68), Gly (68), Ile (65), Met (3), and Trp (67). The amino acids were arranged in a repeating manner within the full-length epsin protein. We are the first to report a full-length protein conists of such a minimum number of essential amino acids. Similarly, the hypothetical protein with the highest *pI* (13.96) was composed of only nine amino acids, namely Ala (62), Gly (132), Lys (127), Leu (197), Met (M), Pro (4), Gln (64), Arg (132), and Ser (66). Intriguingly, cysteine, which is one of the most important amino acids as it is responsible for the formation of disulphide bonds, was not found in either the smallest or largest protein. Disulphide bonds provide the conformational stability of protein and are commonly found in extracellular proteins and only rarely in intracellular proteins [52]. The absence of Cys amino acids in the above-mentioned proteins suggests, they are localized to the intracellular compartments within the cell.

The plant proteome is primarily composed of acidic *pI* proteins rather than basic *pI* proteins (Additional file 2: Table S1). Approximately, 56.44% of the analysed proteins had a *pI* within the acidic *pI* range with an average of *pI* 5.62 (Fig. 1, Additional file 2: Table S1). The average (%) of acidic *pI* proteins was comparatively higher in the lower eukaryotic plants, algae, and bryophytes, than in the higher land plants. A total of 64.18% of proteins in *Chlamydomonas eustigma* were found in the acidic *pI* region, followed by *Ostreococcus lucimarinus* (64.17%), *Micromonas commoda* (63.30%), *Helicosporium* sp. (62.97%), *Gonium pectoral* (62.76%), *Chromochloris zofin*giensis (62.41%), *Coccomyxa subellipsoidea* (62.12%), and *Sphagnum fallax* (61.83%). The algal species, *Porphyra umbilicalis,* had the lowest percentage (29.80%) of acidic *pI* proteins. The dicot plant, *Punica granatum,* and the algal species, *Botrycoccus braunii,* had a significantly lower percentage of acidic *pI* proteins (45.72 and 47.18%, respectively) relative to other plants. Principal component analysis (PCA) of acidic *pI* protein content revealed that the acidic proteins of bryophytes and monocots cluster closely to each other compared to algae and eudicot plants (Fig. 2). Similarly, in the case of basic *pI* proteins, a great variation was observed for algae, eudicot and monocot plants (Fig. 3). The basic *pI* proteins of bryophytes, however, were found to be consistent. A previous study reported that protein *pI* values are correlated with the sub-cellular localization of the proteins, and that the *pI* of cytosolic proteins fall below 7 [21]. Among cytosolic proteins are those involved in 26S proteasome degradation, oxidative pentose phosphate pathway, actin/tubulin, mevalonate pathway, sugar and nucleotide biosynthesis, glycolysis, RNA processing, and several other cellular process. Our analysis indicated that the *pI* of all cytosolic proteins does not fall in the acidic *pI* range. Ribosomal proteins, pre-mRNA splicing factors, transcription factors, auxin induced protein, extensin, senescence associated protein, cyclin dependent protein kinase and other cytoplasmic proteins had a *pI* greater than 7.

**Fig. 2.**
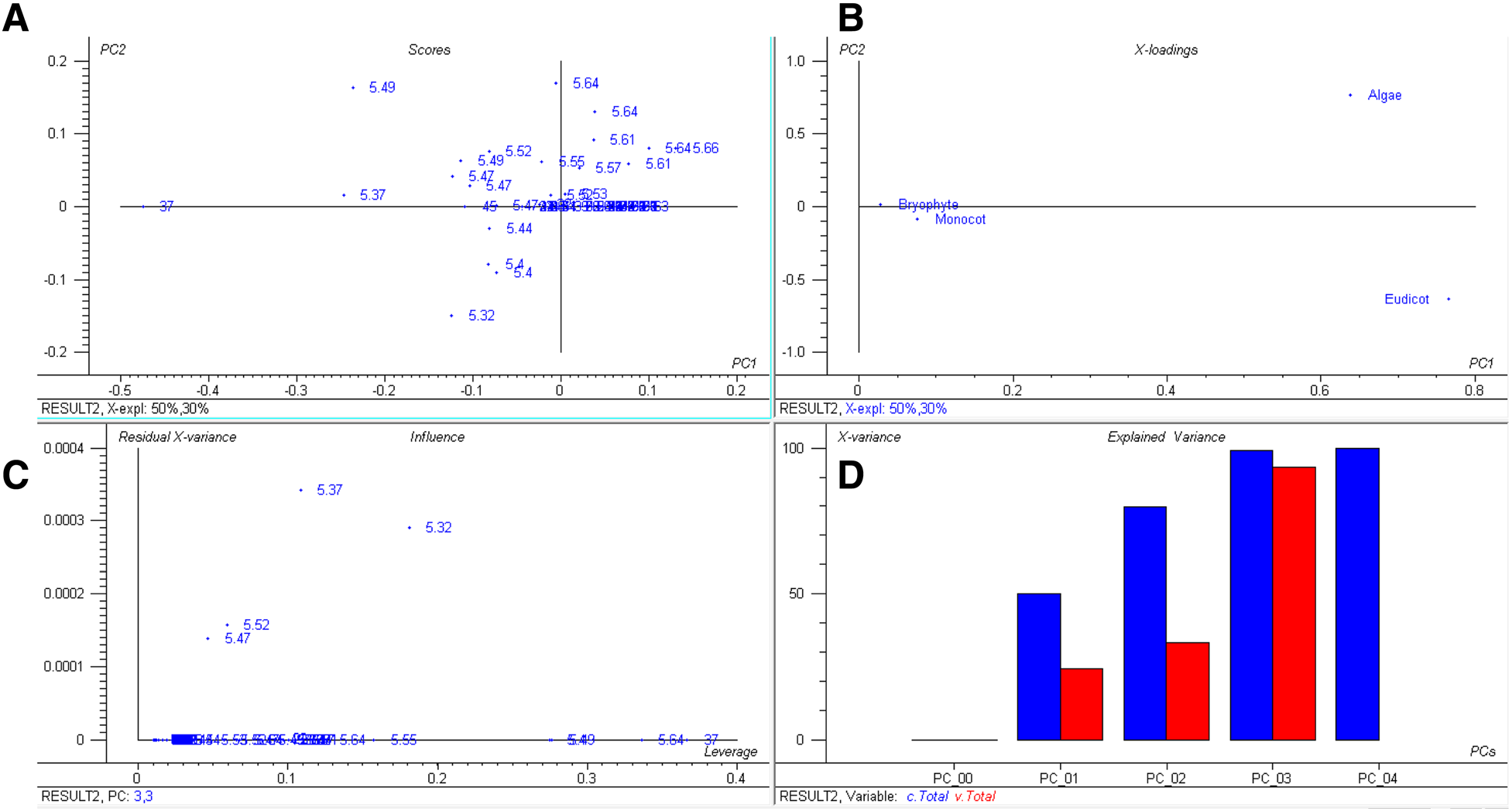
Principal component analysis (PCA) of acidic *pI* proteins. The PCA plot illustrates the relationship between the acidic *pI* of bryophytes and monocot plants which exhibit a linear correlation relative to algae and eudicots. In the figure (**a**) scores: show the similarities in sample grouping, **b** loading: represents the relative position of a variables and how it relates the samples to the different variables (**c**). Influence plot: represents the Qor F-residuals vs. Leverage or Hotelling T2 statistics that show the residual statistics on the ordinate axis of sample distance to model, and (**d**) variance: represents the variation in the data described by the different components. Total residual variance is computed as sum of square of residuals for all the variables, divided by the number of degrees of freedom. The green colour indicates the calibration and the red indicates

**Fig. 3.**
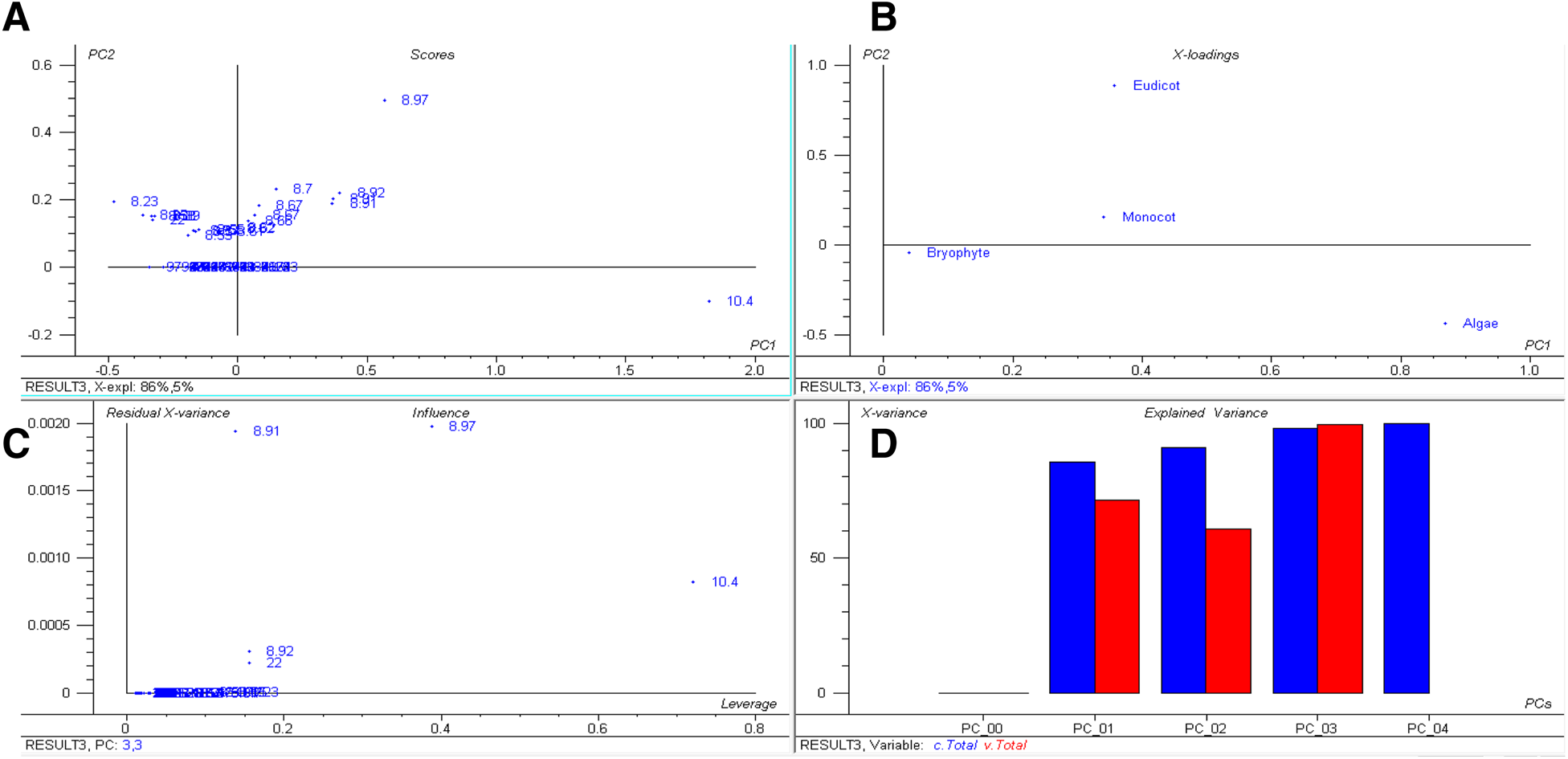
Principal component analysis (PCA) of basic *pI* proteins. The PCA plot illustrates that the basic *pI* of algae, bryophytes, eudicots, and monocot plants cluster distinctly from each other and that there is no lineage-specific correlation with basic *pI* proteins. In the figure (**a**) scores: show the similarities in sample grouping, (**b**) loading: represents the relative position of a variables and how it relates the samples to the different variables (**c**) Influence plot: represents the Qor F-residuals vs. Leverage or Hotelling T2 statistics that show the residual statistics on the ordinate axis of sample distance to model, and (**d**) variance: represents the variation in the data described by the different components. Total residual variance is computed as sum of square of residuals for all the variables, divided by the number of degrees of freedom. The green colour indicates the calibration and the red indicates the validation

In contrast to acidic *pI* proteins, plants possess a comparatively low number of basic *pI* proteins. On average, 43.34% of the analysed plant proteins possessed a *pI* in the basic range with an average pI of 8.37 (Fig. 1, Additional file 2: Table S1). The highest percentage of basic *pI* proteins was found in *Porphyra umbilicalis,* where 70.09% of the proteins had a basic *pI* (Additional file 2: Table S1). *Punica granatum* also had a high percentage (54.11%) of basic *pI* proteins (Additional file 2: Table S1). The lowest percentage of basic *pI* proteins was found in the algal species, *Chlamydomonas eustigma* (35.56%), followed by *Ostreococcus lucimarinus* (35.65%), *Micromonas commoda* (36.52%), *Helicosporidium* sp. (36.89%), and *Gonium pectorale* (37.04%). It is difficult to establish the reason that algal species contain more acidic *pI* and less basic *pI* proteins. *Porphyra umbilicalis* is a cold-water seaweed within the family, Bangiophyceae, and it is the most domesticated marine algae. The 87.7 Mbp haploid genome of *P. umbilicalis* has a 65.8% GC content and an evolutionary study reported that the genome of *Porphyra umbilicalis* had undergone a reduction in size [53]. Since this species is found in the intertidal region of the ocean, it has developed the ability to cope with mid-to-high levels of tidal stress. *Porphyra* is also tolerant to UV-A and UVB radiation [53–55]. The high GC content in *Porphyra umbilicalis* is directly proportional to the high percentage of basic proteins. The GC content of algal species is higher relative to other plant species and algal species possess a lower percentage of basic *pI* proteins. This suggests that, in algae, percentage GC content is inversely proportional to percentage of proteins with a basic *pI*. However, this is not true in the case of higher plants.

### The pI of plant proteomes exhibits a trimodal distribution

The *pI* of the analysed plant proteins ranged from 1.99 to 13.96 and exhibited a trimodal distribution (Fig. 4). Schwartz et al., previously reported a trimodal distribution of the *pI* of eukaryotic proteins [21], however, they did not provide information on the number of sequences/species considered in their study. Proteins are typically soluble near their isoelectric point and the cytoplasm possesses a pH that is close to neutral. This might be the possible reason for the trimodal distribution of *pI*. Although the *pI* values of proteins estimated in silico or experimentally might be different in vivo, they are typically in close agreement and represents a virtual 2-DE gel [56]. However, the acidic peak was more prominent than the alkaline (Fig. 4). Kiraga et al., (2006) reported a bimodal distribution of the *pI* of proteins from all organisms, citing acidic and basic *pI* as the basis of the modality [19], where modality is defined as the set of data values that appears most often. They reported that taxonomy, ecological niche, proteome size, and sub-cellular localization are correlated with acidic and basic proteins. Notwithstanding, no correlation was found in the present study between either acidic or basic *pI* of proteins with regard to taxonomy, ecological niche, or proteome size. For example, *Hordeum vulgare* and *Brassica napus* possess the largest proteomes among the studied plant species, possessing 248180 and 123465 proteins, respectively. In *H. vulgare,* 53.28% of the proteins fall in the acidic and 46.50% fall in the basic *pI* ranges; while in *B. napus*, 55.28% of the proteins have an acidic *pI* and 44.48% have a basic *pI*. Other species with smaller proteomes, however, possess a higher percentage of acidic or basic proteins (Additional file 2:Table S1). Therefore, no correlation exists between the percentage of either acidic or basic proteins and proteome size, taxonomy, or the ecological niche of an organism. Knight et al. also addressed a negative correlation between the *pI* of a protein and phylogeny of the organism [57]. The presence of a trimodal distribution of the *pI* of the plant proteome can be considered as a virtual 2D-gel of a plant’s proteins where the *pI* of the protein is plotted against the molecular weight of the protein. On average, 0.21% of the analysed proteins were found to have a neutral *pI* (*pI* 7), while only 0.09% of the proteins in *O. lucimarinus* fall in neutral *pI*. The modal distribution of protein *pI* shows its physiochemical properties rather than sequence evolution [58].

**Fig. 4.**
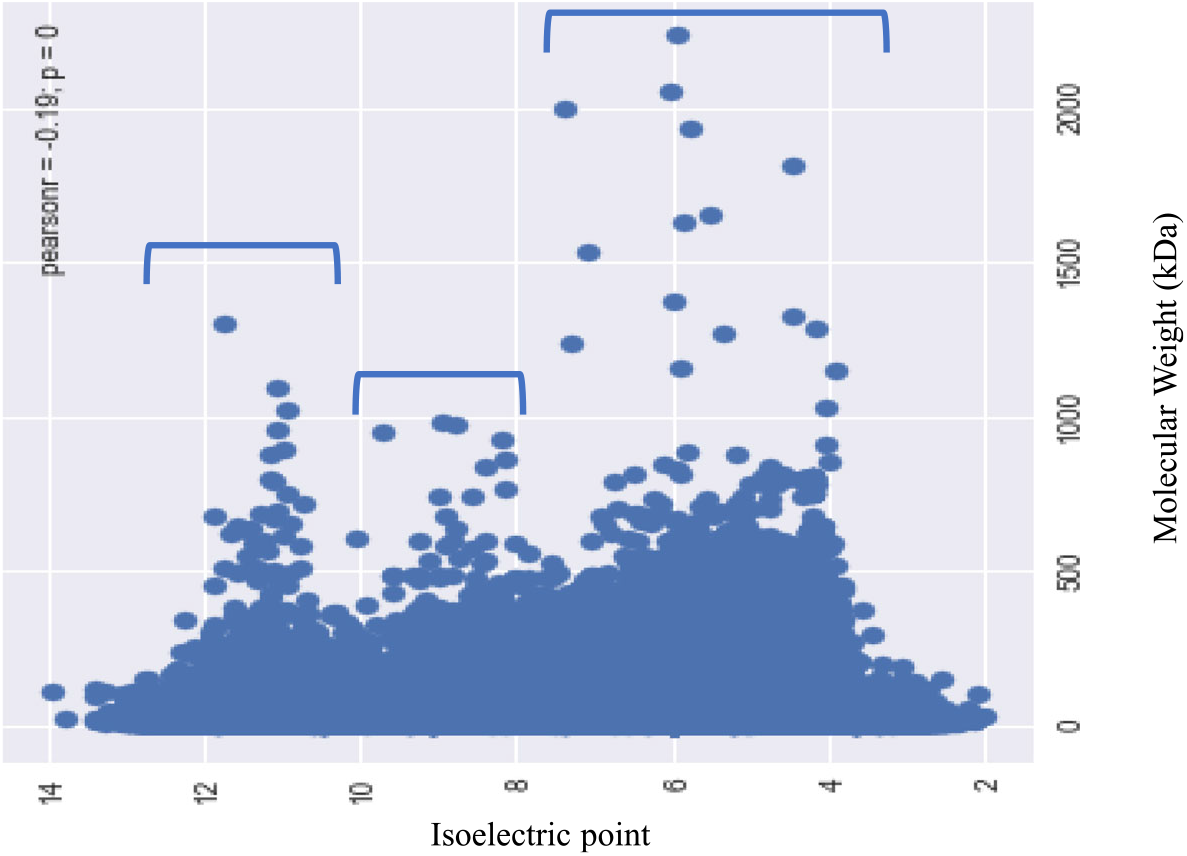
Trimodal distribution of isoelectric points (*pI*) and the molecular mass (kDa) of plant proteins. The *pI* of plant proteins ranged from 1.99 (epsin) to 13.96 (hypothetical protein), while the molecular mass ranged from 0.54 (unknown) to 2236.8 (type I polyketide synthase) kDa. The Xaxis represents the *pI* and the Yaxis represents the molecular mass of the proteins

### Leu is a highand Trp is a low-abundant amino acid in the plant proteome

The plant-kingdom-wide proteome analysis of amino acid composition revealed that the Leu was the most (9.62%) while Trp was the least (1.28%) abundant amino acid (Fig. 5, Additional file 1). Leu is a nonpolar amino acid, whereas Trp contains an aromatic ring. The distribution of amino acids indicates that the synthesis of nonpolar amino acids is more favoured in the plant proteomes than the polar amino acids or those containing an aromatic ring. The average abundance of other nonpolar amino acids Ala, Gly, Ile, Met, and Val was 6.68, 6.80, 4.94, 2.40, and 6.55%, respectively (3 Table 2, Additional file 1). Trp and Tyr amino acid contain an aromatic ring and the abundance of these two proteins is relatively low in the plant proteome compared to other amino acids. Results of the conducted analysis indicated that the abundance of Ala (17.58%), Gly (11.76%), Pro (9.2%), and Arg (9.81%) were the highest; whereas, Tyr (1.33%), Gln (2.04%), Asn (1.53%), Met (1.45%), Lys (7.07%), Lys (2.08%), Ile (1.77%), Phe (2.01%), and Glu (3.52%) were the lowest in *Porphyra umbilicalis*. In a few algae and seaweeds Ala, Asp, Glu, Gly, Pro, Gln, Arg, Thr, and Val were found in high percentage while Asp, Glu, Phe, His, Ile, Lys, Leu, Met, Asn, Gln, and Ser were found in low percentage (Additional file 3: Table S2). These findings indicate that the composition of amino acids in unicellular algae, seaweeds, and gymnosperms are more dynamic and variable than in angiosperms and other terrestrial land plants. Principal component analysis revealed that the low abundant amino acids, Trp, Tyr, His, Met, Cys, and Xaa (unknown), cluster in one group while the high abundant amino acids, Leu, Glu, Ile, Lys, and Ser, cluster in another group (Fig. 6). None of the terrestrial land plants were located in the high- and low-abundant amino acid clusters. This put forward the fact that the amino acid composition and proteome of the land plants are more conserved and stable relative to the algae and seaweeds. In addition, the PCA analysis revealed that the *pI* of algae, eudicots, and monocots are lineage specific. The *pI* of algae, monocots, and eudicots were strongly correlated and clustered together (Fig. 6).

**Fig. 5.**
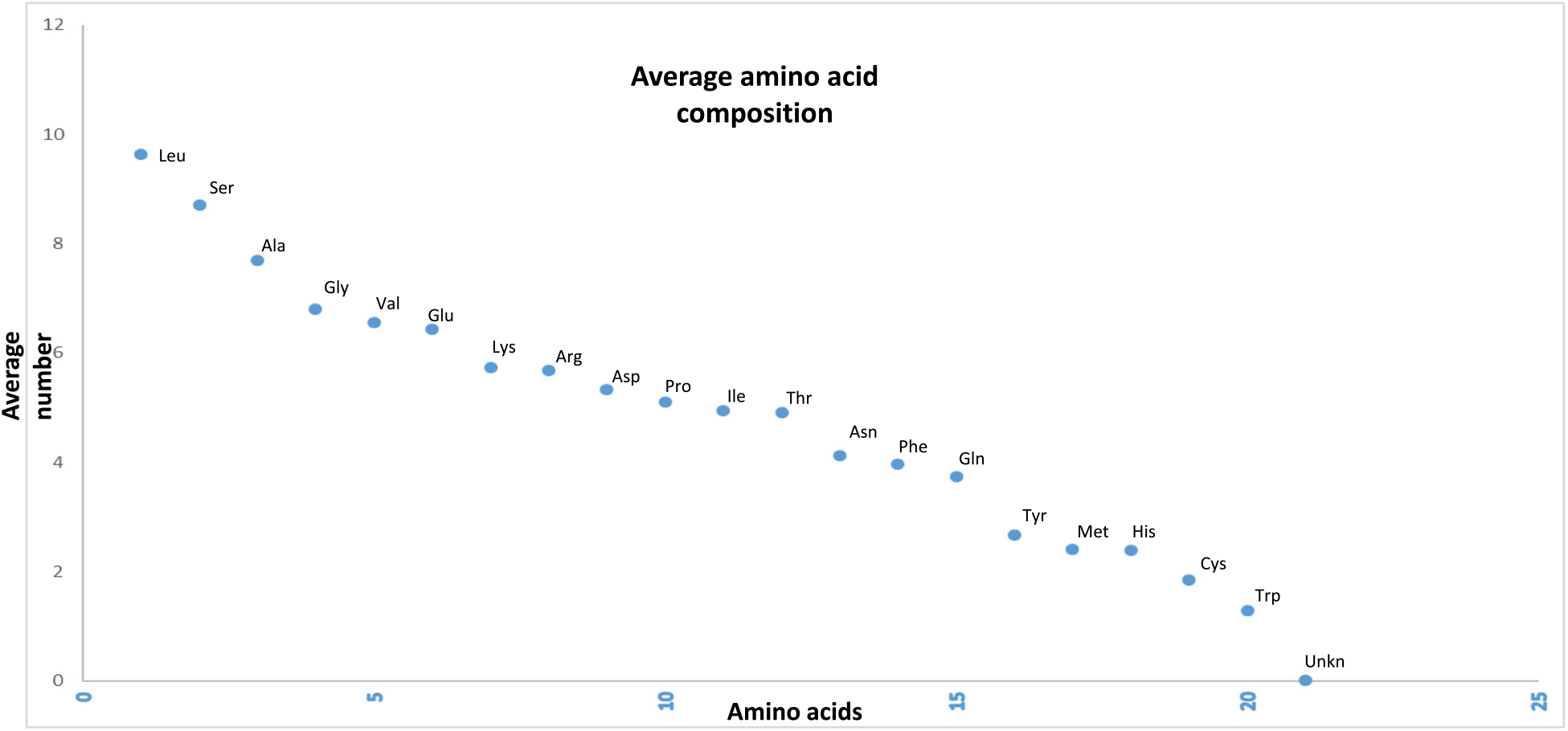
Average amino acid composition of proteins in the plant kingdom. Leu is the most abundant while Trp is the least abundant. The amino acid, Sec, was only found in a few species of algae and was absent from all other species. Ambiguous amino acids were found in a few species as well

**Fig. 6.**
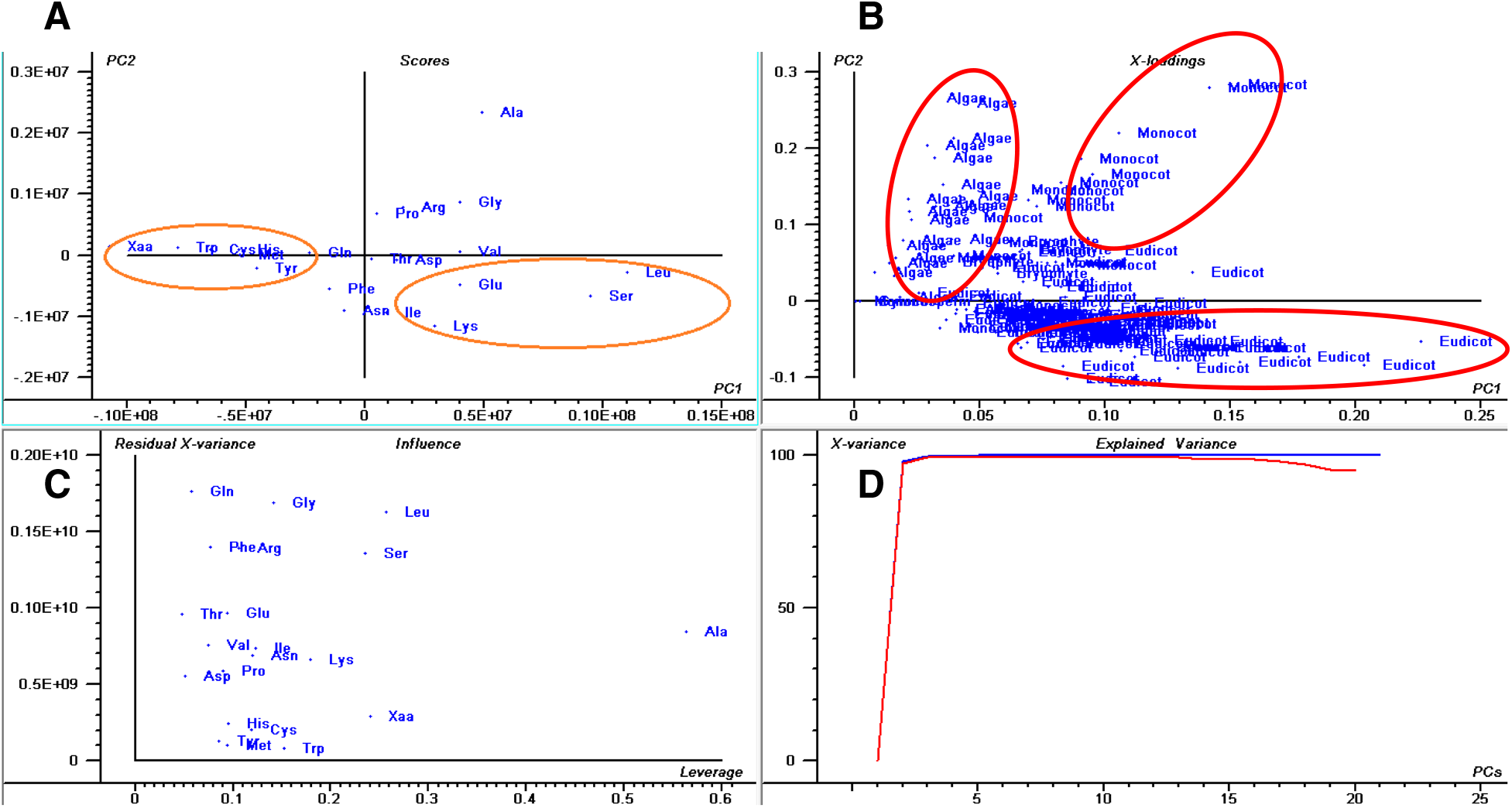
Principal component analysis (PCA) of amino acid abundance in plant proteomes. The PCA plot shows that Tyr, Trp, Cys, His, Met, and Xaa (unknown) amino acids are low-abundance and cluster together. The abundance of Leu, Ser, Ile, Lys, and Gln was higher and grouped together. The plot shows that the abundance of amino acids is lineage specific. Algae, eudicots, and monocot plants exhibit a lineage specific correlation. In the figure (**a**) scores: show the similarities in sample grouping, (**b**) loading: represents the relative position of a variables and how it relates the samples to the different variables (**c**) Influence plot: represents the Qor F-residuals vs. Leverage or Hotelling T2 statistics that show the residual statistics on the ordinate axis of sample distance to model, and (**d**) variance: represents the variation in the data described by the different components. Total residual variance is computed as sum of square of residuals for all the variables, divided by the number of degrees of freedom. The green colour indicates the calibration and the red indicates the validation

The question arises, however, as to why the plant proteome contains the highest percentage of Leu and of the lowest percentage of Trp amino acids. Do the energy requirements of the different biosynthetic pathways play a pivotal role in deciding the abundance of amino acids in a proteome? To address this question, an study was conducted to figure out the role of amino acid biosynthetic pathways in determining the abundance of specific amino acids in the proteome. The amino acid composition are used to deduced the properties of proteins that are selected in evolution for their chemical properties [59]. Most importantly, longer protein helps to form complex secondary and tertiary structure. In addition, amino acid composition influence the core and surface protein structure.

Various amino acids are produced in different biosynthetic pathways [60–64] (Fig. 7). In few cases, some of the amino acids act as the substrate for the biosynthesis of other amino acids; whereas in other cases, allosteric inhibition of the biosynthesis of amino acids occurs [65–67]. In all of these amino acid biosynthetic pathways, along with substrate, ATP or NADH/NADPH are used as a source of energy. Overall, the biosynthesis of 20 essential amino acid families are grouped by metabolic precursors [68] (Table 1); namely α-ketoglutarate (Arg, Gln, Glu, Pro), pyruvate (Ala, Ile, Leu, Val), 3-phosphoglycerate (Cys, Gly, Ser), phosphoenolpyruvate and erythrose 4-phosphate (Phe, Trp, Tyr), oxaloacetate (Asn, Asp, Lys, Met, Thr), and ribose 5-phosphate (His) (Table 1) [68]. Ile, Alan, Leu, and Val are synthesized from pyruvate; Glu, Arg, Gln, and Pro are synthesized from α-ketoglutarate whereas Gly and Ser are synthesized from 3-phosphoglycerate [68]. The abundance of these amino acids in plant proteomes are considerably higher relative to the other amino acids (Fig. 7, Table 3). 3-phosphoglycerate and pyruvate are act as transient product of glycolysis and the amino acids produced from these intermediates maintain a high abundance in the plant proteome. The intermediate, 3-phosphoglycerate, is formed in an early step of glycolysis [68]. Gly and Ser are produced from 3-phosphoglycerate and quite bountiful in the plant proteome (Fig. 7, Table 3). The amino acid Cys, which is also synthesized from 3-phosphoglycerate [68], however, is present in low abundance (1.85%) in the plant proteome. The low richness of Cys possibly due to the allosteric inhibition. Phe (3.97), Trp (1.28%), and Try (2.67%) contain an aromatic ring and are synthesized via phosphoenolpyruvate and erythrose 4-phospahte. The aromatic amino acids including Trp, Tyr, and Phe are in low abundance in the plant proteome (Table 1). Since Phe also plays a role in the biosynthesis of Tyr, the abundance of Phe is relatively higher than Trp and Tyr. Glucose 6-phosphate gives rise to ribose 5-phosphate in a complex reaction of four steps [68] and His gets subsequently synthesized from ribose 5-phosphate. It is possible that the complexity of the biosynthetic pathways of amino acids containing ring compounds might be the reason for their low abundance in the plant proteome.

**Table 1.**
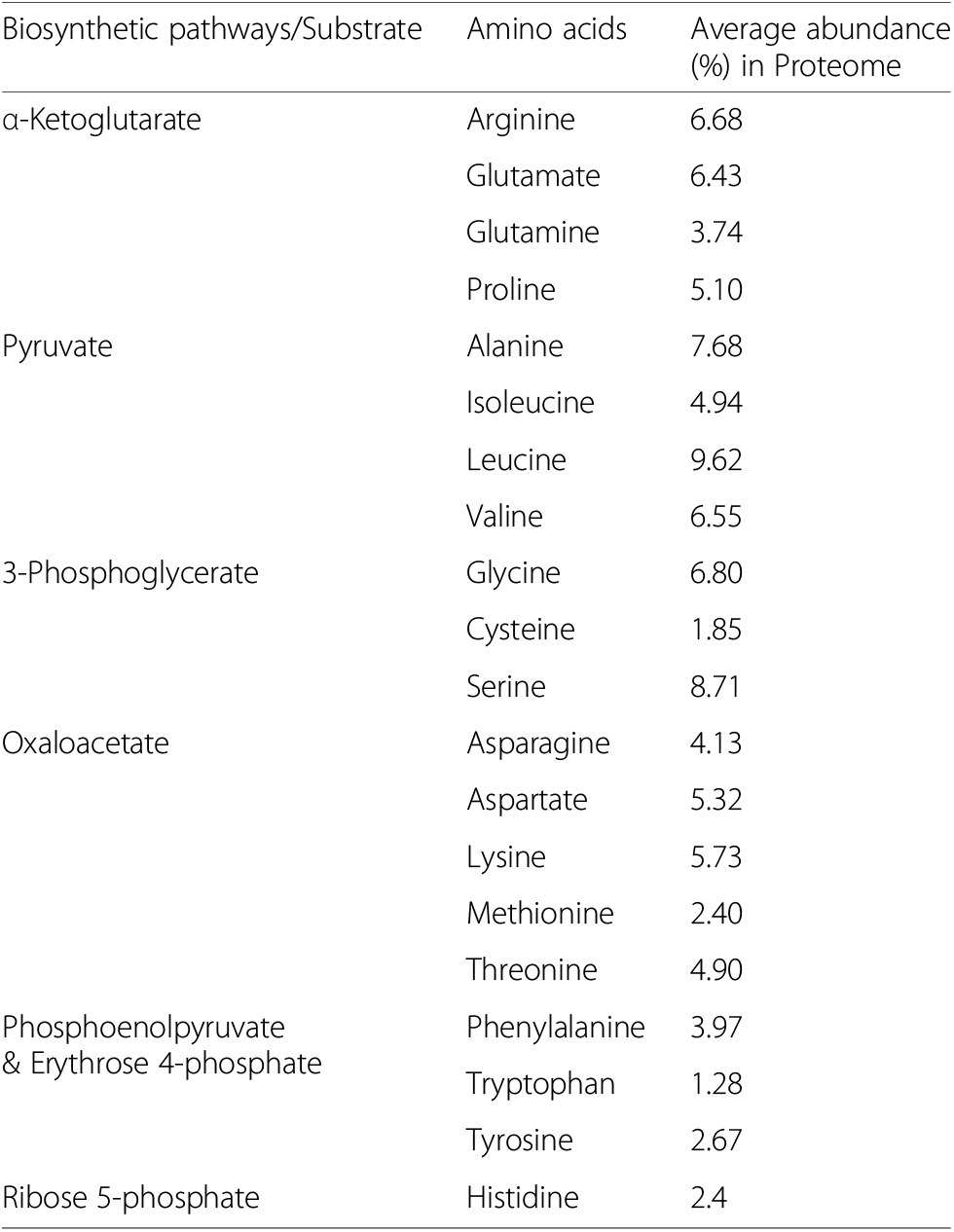
Average abundance of different amino acids in plant proteome. Leu was high abundant whereas Trp was the low abundant amino acid in the plant kingdom. The average amino acid composition includes 5.8 million protein sequences from 145 plant species

**Fig. 7.**
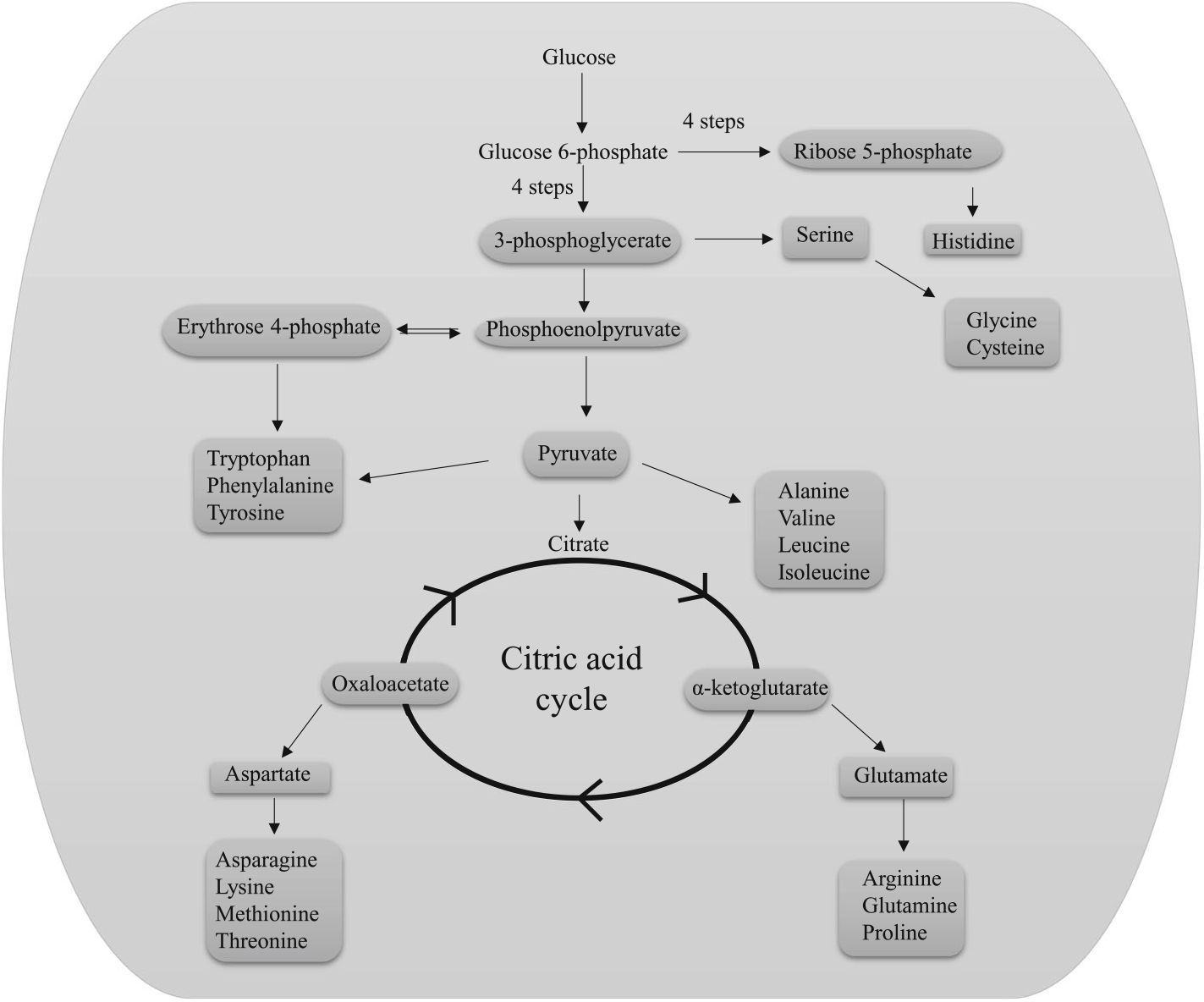
Schematic illustration of the biosynthetic pathway of amino acids. The abundance of aromatic ring containing amino acids is lower relative to other amino acids. The average abundance of the aromatic ring containing amino acid, Trp, is the lowest amongst others that are biosynthesized via phosphoenolpyruvate and erythrose 4-phosphate. Similarly, the abundance of Cys is relatively low compared to other amino acids. Ser is biosynthesized from 3-phosphoglycerate and Ser is subsequently used to produce Gly and Cys amino acids. The abundance of Cys is lower relative to Gly, suggesting the existence of allosteric feed-back inhibition of the biosynthesis of Cys by Ser

### Plants possess selenocysteine (sec) and other novel amino acids

A few of the plant proteomes that were analysed had proteins containing the amino acid, selenocysteine (Sec) encoded by TGA (opal) codon [69]. *C. reinhardtii*, *M. pusilla*, and *V. carteri* contained 9, 16, and 11 Sec amino acids in their proteome, respectively. Selenium containing proteins are frequently found in the proteome of bacterial and animal kingdom [69–71]. However, it has been reported to be present in the plant species as well. Novoselov et al., (2002) reported the presence of a selenoprotein in *C. reinhardtii* [72]. In our analysis, nine selenoproteins (selenoprotein H, selenoprotein K1, selenoprotein M2, selenoprotein T, selenoprotein U, selenoprotein W1, selenoprotein W2, NADPH-dependent thioredoxin reductase 1, and glutathione peroxidase) were identified in *C. reinhardtii*. In addition, *M. pusilla* was found to possess 14 Sec-containing proteins [DSBA oxidoreductase (2 no.), glutathione peroxidase (4 no.), selenoprotein T, selenoprotein W, selenoprotein, selenoprotein M, selenoprotein H, selenoprotein O, methyltransferase, and peroxiredoxin). In addition, *V. carteri* was found to possess 10 Sec containing proteins (selenoprotein T, selenoprotein K1, selenoprotein H, selenoprotein W1, selenoprotein M2, selenoprotein U, glutathione peroxidase, membrane selenoprotein, NADPH-dependent thioredoxin reductase, and peptide methionine-S-sulfoxide reductase). According to our observation, this is the first report regarding the presence of selenoproteins in *M. pusilla* and *V. carteri* and the first to report the presence of H, K1, T, U, M, M2, O, W1, and W2 selenoprotein families in *C. reinhardtii*, *M. pusilla*, and *V. carteri*. The I, N, P, R, S, and V selenoprotein family members are commonly found in the animal kingdom [73] but are absent in *C. reinhardtii, M. pusilla*, and *V. carteri*. This is also the first report of the Sec-containing proteins, DSBA oxidoreductase, methyltransferase, peroxiredoxin, peptide methionine-Ssulfoxide reductase, and membrane selenoprotein in plants lineage (algae). Notably, the selenoproteins DSBA oxidoreductase, methyltransferase, peroxiredoxin, peptide methionine-S-sulfoxide reductase, and membrane selenoprotein have not been reported in animal species. Outside of algal species, no other plant species; including bryophytes, pteridophytes gymnosperms and angiosperms, were found to possess a selenoprotein. Selenoprotein is superimposed in the translation machinery and more specifically selenoprotein mRNA contain selenocysteine insertion sequence element (SECIS) downstream to the Sec-encoding UGA codon [74, 75]. The SECIS-sequence recognise and binds the selenocysteine-specific elongation factor (*selB*) and forms complex with tRNA-^Sec^ (*selC*). The tRNA^Sec^ acylated with serine by seryl-tRNA synthastase and subsequently transformed to Sec-tRNA^Sec^ by Sec synthase (*selA*). Later, selA utilizes selenophosphate as the selenium donor that synthesized by selenophosphate synthetase (selD) [71, 75]. The selenium is found in the active site of selenoproteins, more specifically involved in redox reactions [71]. In human, selenoprotein H regulates cell cycle progression and proliferation of human colorectal cancer cells [76]. Selenoprotein K enhances protein palmitoylation [77, 78], selenoprotein T attenuates the development of left ventricular dysfunction after myocardial infraction in rats [79].

Some plant proteomes were also found to possess a few unknown or unspecified amino acids, commonly designated as Xaa (X). Among the analysed plant species, *Aegilops tauschii, Amaranthus hypochondriacus*, and *Amborella trichocarpa* encoded 149377, 55412, and 25843 X amino acids, respectively. *Solanum lycopersicum* was found to contain only one X amino acid, while at least 18 species (*Solanum pennellii, Solanum tuberosum, Sorghum bicolor, Sphagnum fallax, Spinacia oleracea, Spirodela polyrhiza, Tarenaya hassleriana, Theobroma cacao, Trifolium pratense, Trifolium subterraneum, Triticum aestivum, Triticum urartu, Vigna angularis, Vigna radiata, Vigna anguiculata, Vitis vinifera, Volvox carteri,* and *Zostera marina*) were found to lack any Xaa amino acids in their proteome. Xaa amino acids are commonly considered as non-protein amino acids as they are yet to be reported as protein coding amino acids. Among the studied plant species, ten were found to contain amino acid B (Asx) that codes for the ambiguous amino acid Asn or Asp that is translated as Asp. Species that were found to possess an Asx amino acid included *Arachis duranensis* (1), *Brachypodium stacei* (40), *Dichanthelium oligosanthes* (20), *Dunaliella salina* (31), *Glycine max* (1), *Malus domestica* (4080), *Momordica charantia* (98), *Nelumbo nucifera* (64), *Prunus persica* (1), and *Trifolium pratense* (76). At least six species were found to possess a J (Xle) amino acid. Xle amino acid can encode either Leu or Ile but during translation produces Leu. Species that were found to possess Xle amino acids included *Arabidopsis thaliana* (10), *Dichanthelium oligosanthes* (11), *Malus domestica* (2175), *Momordica charantia* (39), *Nelumbo nucifera* (29), and *Trifolium pratense* (39). At least seven species were found to possess a Z (Glx) amino acid that codes for either Glu or Gln, which is subsequently translated as Glu. Species that were found to encode a Glx amino acid included *Brachypodium stacei* (20), *Dichanthelium oligosanthes* (16), *Dunaliella salina* (7), *Malus domestica* (1552), *Momordica charantia* (28), *Nelumbo nucifera* (14), and *Trifolium pratense* (25). Among the studied species, *Malus domestica* was found to contain highest number of ambiguous amino acids (Asx, Xle, and Glx). Bodley and Davie (1966) reported the incorporation of ambiguous amino acids in a peptide chain [80]. The presence of ethanol or streptomycin or a high magnesium ion concentration induces ambiguous coding in the peptide chain [80]. They reported that poly-U (uridylic acid) in the presence of a high concentration of magnesium ions or ethanol or streptomycin induces the incorporation of Leu/Ile amino acids in a peptide chain [80]. This explains how the specificity of the protein translation process can be altered by the presence of environmental factors. A high concentration of magnesium ions, organic solvents, antibiotics, pH, and low temperature have the ability to modify the coding specificity of a peptide chain [80]. Under some conditions poly-U triggers the incorporation of Leu and Ile or Phe [80]. *Malus domestica* is rich in magnesium ions (1%) and this might explain the presence of such a high number of ambiguous amino acids in its proteome. The presence of D-Xaa-D-Xaa-tRNA in the P-site lead to ribosomal stalling of elongation factor G (EF-G) [81].

## Conclusion

A proteome-wide analysis of the plant kingdom identified proteins with a great range of molecular mass and isoelectric points. Isoelectric points ranged from 1.99 to 13.96, covering almost the entire pH range. It is quite intriguing to think about the functions of protein at *pI* 1.99 or 13.96. Proteins with an acidic *pI* predominate over the proteins with an alkaline *pI*, and the presence of proteins with a *pI* that is near neutral is very negligible. The percentage of proteins with acidic or basic *pI* is not related to the host cell, alkalinity or acidity of the environment. The trimodal distribution of *pI* of plant proteome is due to the differential composition of amino acids in different species and this might be associated with the environmental and ecological pressure. Rate of mutation and nucleotide substitution can also be the other responsible factors associated with composition of the amino acids and hence the *pI*. Additionally, the GC content of a genome or size distribution of the overall proteome are not directly proportional to the distribution (percent basic vs. percent acidic) of the *pI* in the proteome. The presence of Sec-containing proteins in some algal species needs to be further investigated to determine their functional role. Similarly, the presence of ambiguous amino acids in plant species should be further evaluated individually. The presence of a pyrrolysine amino acid in the plant kingdom was not observed in the present study.

## Methods

Protein sequences of the entire proteome of the analysed plant species were downloaded from the National Center for Biotechnology Information (NCBI) and Phytozome, DOE Joint Genome Institute (https://phytozome.jgi.doe.gov/pz/portal.html). All of the studied sequences were annotated nuclear-encoded proteins. The isoelectric point of each protein of each of the analysed plant species was calculated individually using the Python-based command line “Protein isoelectric point calculator” (IPC Python) in a Linux platform [2]. The source code used was as written by Kozlowski (2016).

Once the molecular mass and isoelectric point of the proteins in each species was determined, they were separated into acidic and basic *pI* categories. Subsequently, the average *pI* and percentage of proteins in each category was calculated using a Microsoft excel worksheet. A graph comparison of isoelectric point versus molecular mass was prepared using a python-based platform. Pearson-correlation (r = 0.19, *p* = 0) was used for the association analysis of molecular mass and isoelectric point. The X-axis data statistics were as follows: mean, 4.717365e+ 01; std., 3.662983e+ 01; min, 8.909000e-02; 25%, 2.279452e+ 01; 50%, 3.874486e+ 01; 75%, 5.999628e+ 01, and max, 2.236803e+ 03. The Y-axis data statistics were: mean, 6.840657e+ 00; std., 1.594912e+ 00; min, 1.990000e+ 00; 25%, 5.537000e+00; 50%, 6.605000e+ 00; 75%, 8.053000e+ 00, and max, 1.396300e+ 01.

### Principal component analysis

Principal component analysis of the plant proteome parameters was carried out using a portable Unscrambler software version 9.7 using the excel file format. For acidic and basic *pI*, the plant proteome data were grouped according to the plant lineage algae, bryophyte, monocot, and eudicot plants. The average of acidic and basic *pI* was used to construct the PCA plot. Similarly, amino acid abundance was also analysed in relation to algae, bryophyte, eudicot, and monocot lineage.

## Additional files

### Additional file 1

Supplementary file containing average amino acid composition of plant proteomes. The file is present in excel sheet and individual sheet represents details of different amino acid. (XLSX 252 kb)

### Additional file 2

**Table S1.** Details of plant proteome. Table shows acidic pI of proteins predominates the basic *pI*. However, in sea weed *Porphyra umbilicalis*, basic *pI* predominates over the acidic *pI*. Putative polyketide synthase type I found in lower eukaryote *Volvox carteri* was found to be the largest protein in the plant lineage. However, titin was found to be the largest protein in the higher eukaryotic land plants. Asterisks represents no specific data available for the said item. (DOCX 61 kb)

### Additional file 3

**Table S2.** Abundance of various amino acids in different species. The second column represents the average abundance whereas the third and fourth column represent variation (high and low) in amino acid composition in different species from the average. (DOCX 16 kb)

## Abbreviations

60S RP L41: 60S ribosomal protein subunit L41
AAPT: Aminoalcoholphosphotransferase
ATF7IP: Activating transcription factor 7-interacting protein 1
CDPK: Cyclin-dependent serine/threonineprotein kinase
CSF: Cleavage stimulation factor subunit 2 tau variant-like
CWP: Cell wall protein
DDG: Dolichyl-diphosphooligosaccharideprotein glycosyltransferase subunit 4A
DDGT: Dolichyldiphosphooligosaccharideprotein glycosyltransferase subunit 4A
DDHGT 4A: Dolichyl-diphosphooligosaccharideprotein glycosyltransferase subunit 4A
ER TF: ethylene responsive transcription factor
FS CAYBR BP: Fibrous sheath CABYR-binding protein-like
GRCW protein: Glycine-rich cell wall structural protein
IMP: inosine-5′-Monophosphate cyclohydrolase
LEA: Late embryogenesis associated protein
NFD: Nuclear fusion defective
PERK2: Proline-rich receptor-like protein kinase PERK2
PKDP: Polycystic kidney disease protein 1-like 3
PKS: Polyketide synthase
PPCD VHS3-like: Phosphopantothenoylcysteine decarboxylase subunit VHS3-like isoform X2
PPCD: Phosphopantothenoylcysteine decarboxylase
PS-I RC N: Photosystem I reaction center subunit N, chloroplastic, partial
RBP: RNA binding protein
RNA Pol. II Med 17: Mediator of RNA polymerase II subunit 17
RPB1: DNA-directed RNA polymerase II subunit RPB1-like isoform X3
SARMP: Serine/arginine repetitive matrix protein 2-like
SCP: Spore coat protein
SR45: Serine/arginine-rich splicing factor SR45-like
SVC: Satellite virus coat protein
ZRF: Zinc ring finger-type

## Acknowledgements

Not available.

## Availability of data materials

All the studied data were taken from publicly available databases and data associated with the manuscript is provided in supplementary file.

## Competing of interest

The authors declare that they have no competing interests.

## Authors’ contributions

TKM: conceived the idea, collected the protein sequences, analysed and interpreted the data, drafted the manuscript, ALK: revised the manuscript, AH: drafted and revised the manuscript, EFA: drafted and revised the manuscript, AAH: revised the manuscript. All authors read and approved the final manuscript.

## Funding

Not available.

## Ethics approval and consent to participate

Not applicable.

## Consent for publication

All authors agree and have consent for publication.

## Publisher’s Note

Springer Nature remains neutral with regard to jurisdictional claims in published maps and institutional affiliations.

